# Striking clone-specific nymph deformity in pea aphid exposed to heat stress

**DOI:** 10.1101/2023.02.15.528785

**Authors:** Hawa Jahan, Mouhammad Shadi Khudr, Ali Arafeh, Reinmar Hager

## Abstract

Climatic changes, such as heatwaves, pose unprecedented challenges for insects, as escalated temperatures above the thermal optimum alter insect reproductive strategies and energy metabolism. While thermal stress responses have been reported in different insect species, thermo-induced developmental abnormalities in phloem-feeding pests are largely unknown. In this laboratory study, we raised two groups of first instar nymphs belonging to two clones of the pea aphid *Acyrthosiphon pisum*, on fava beans *Vicia faba*. The instars developed and then asexually reproduced under constant exposure to a sub-lethal heatwave (27°C) for 14 days. Most mothers survived but their progenies showed abnormalities, as stillbirths and appendageless or weak nymphs with folded appendages were delivered. Clone N116 produced more deceased and appendageless embryos, contrary to N127, which produced fewer dead and more malformed premature embryos. Interestingly, the expression of the HSP70 and HSP83 genes differed in mothers between the clones. Moreover, noticeable changes in metabolism, *e.g*., lipids, were also detected and that differed in response to stress. Deformed offspring production after heat exposure may be due to heat injury and differential HSP gene expression, but may also be indicative of a conflict between maternal and offspring fitness. Reproductive altruism might have occurred to ensure some of the genetically identical daughters survive. This is because maintaining homeostasis and complete embryogenesis could not be simultaneously fulfilled due to the high costs of stress. Our findings shine new light on pea aphid responses to heatwaves and merit further examination across different lineages and species.

**Highlights:**

- Parthenogenetic aphids were resilient under heatwave yet produced deformed progeny
- Developmental deformities by clones imply conflicting maternal/offspring fitness
- Clone N127 produced fewer dead and more malformed premature embryos than clone N116
- Expression of two heat shock protein genes differed across clones under heat stress
- Exposure to heat stress led to metabolic differences in the exposed aphids

## Introduction

Global climatic changes, primarily owing to anthropogenic effects [1], have recently been associated with heatwaves that may unprecedentedly occur sooner in early summer in Europe [2–5]. Exposure to high temperatures has complex effects on the metabolism and life-history traits, including the development, fecundity, and population dynamics of herbivorous insects [1,6,7]. The negative impact of exposure to heatwaves may extend to result in serious developmental delay, reproductive malfunctioning, mortality, and even extinction of the insect population exposed to temperatures above optimum thresholds [1,8–10].

Aphids, small soft-bodied phloem-feeding insects, show species- and clone-specific thermal tolerance to abrupt changes in temperature [11]. Zhang et al. (2019) exposed five aphid species, including two clones of *Acyrthosiphon pisum* (Harris), to sub-lethal temperature (38°C) and found distinct differences in their survivability, development time, and fecundity, with higher clone-specific mortality and thermotolerance reported in the exposed pea aphids [12]. Since aphid development, dispersal and reproduction are largely influenced by their sensitivity to thermal changes [13], maintaining a balance between survival, growth, and reproduction is quite challenging under stress and may lead to compromised fitness [14,15].

*Acyrthosiphon pisum*, an important crop pest [16] and eco-evo model organism [17], shows impressive phenotypic plasticity, a phenomenon of producing variable phenotypes in response to environmental stress [18,19]. Reproductive, ontogenetic, and phenotypic plasticities are contextual and may differ across polymorphic lineages, including green and pink morphs, underlined by metabolic differences [21–25]. The optimal temperatures for *A. pisum* range from 20°C to 25°C, with upper limits up to 30°C, depending on the geographic location and adaptability of aphid lineages [13]. Transgenerational effects following exposure to heat stress have been shown to carry over to subsequent generations further affecting offspring development and reproduction [26,27].

It is noteworthy that malnutrition may lead to depressed embryo development and embryo reabsorption [15,28], while embryo retention may also occur as a cost of increased densities in certain secondary endosymbiont communities in aged aphids [29]. However, these phenomena have not been reported in heat-stressed aphids. Thermal stress induces upregulation of heat shock protein genes (*e.g*., Enders et al. 2015 [15]). Particularly, heat shock protein families HSP90 (including HSP83) and HSP70 are associated with repairing denatured proteins and maintaining homeostasis [30]. Impaired production of the HSP genes as well as histone acetyltransferase (HAT) p300/CBP, which is a transcriptional co-regulator, can cause serious embryonic defects [31] and developmental abnormalities [32] in *Drosophila melanogaster* (Meigen) and *A. pisum* [33,34].

Furthermore, Fourier Transform InfraRed spectroscopy (FTIR) is a user-friendly and affordable technique that is utilisable in measuring and understanding the metabolic changes that underpin organism responses to environmental stress [35,36]. FTIR is helpful for metabolomic profiling [24] and structural analysis [37] of insects and their secretions [38], but the application of FTIR to examine thermal stress responses in aphids is still understudied.

In this exploratory laboratory study, we reared two pea aphid clones on *Vicia faba* var *minor* Harz) under thermal optimum (22°C) and exposed them to sub-lethal thermal stress (27°C), resembling a heatwave, for 14 days. Respective aphid first instars developed in each thermal regime until maturity and produced parthenogenetic offspring. We test the following hypotheses: 1) Survival of developing aphids is not affected by exposure to thermal stress with no signs of embryonic anomalies. 2) Aphid clones show universal phenotypic and molecular responses to thermal stress.

## Materials and Methods

### Experimental setup

We established populations of two pea aphid clones (N116 [green] and N127 [pink]) from respective single parthenogenetic apterae. These aphids descended from samples provided by Imperial College London. The clones are of different biotypes, with N116 being more prolific than N127 [39]. The stock aphids were raised on two-week-old fava bean plants (*Vicia faba* var. *minor* Harz) germinated and grown in a growth cabinet at 22°C, 70% RH, and 16D: 8N cycle. The aphid clones were maintained for hundreds of generations in these conditions. We used individual meshed enclosures fitted with plastic pots (9cm x 9cm) filled with compost (Levington F2).

We applied two experimental conditions in terms of temperature (i) thermal optimum (22°C) which was the control as described for the stock culture above, and (ii) heatwave (thermal stress) as a constant high temperature (27°C), based on a pre-experiment pilot. The latter revealed that at 35°C, plants started wilting after a few days of their germination before aphid introduction. At 30°C, both survived plants and the aphids of both clones died within a few days after aphid introduction. At 28°C, aphids survived but barely reproduced. Eventually, at 27°C, both aphid clones survived and reproduced after the introduction of the respective clones, but only a few offspring remained alive after their emergence. We observed and counted unexpected premature or deformed instar nymphs, which was unusual and never reported before out of the aphid mother body.

In the experiment, we always used seven first instars for plant infestation and the instars were introduced to the plant two weeks following germination. We watered the enclosures every two days. There were 2 aphid clones X 2 environmental conditions X 15 replicates = 60 enclosures, see (S1 Fig) for experimental design. The aphids, which survived the heat exposure, matured, and produced offspring, were sampled across the clones and the conditions at the end of the experiment and preserved at −80°C for further assays to examine the expression pattern of HSP genes and to understand whether the aphid clones may show different metabolic fingerprinting in response to the heatwave.

### Statistical analysis

All statistical analyses were done in R ver. 4.0.4 [40]. First, using an Anova model (Type II), *car* package [41], we tested the total number of survived mothers in the microcosm on Day 14 (SM) as a function of (i) Thermal stress (two levels: 22°C [thermal optimum, control], 27°C [thermal stress]); (ii) Aphid clone (N116 [green], N127 [pink]), (iii) plant total dry weight (TDW) as a covariate, and (iv) the interactions of these predictors. Second, we counted the deformed and live nymphs on Day 14. Using a Manova model (Type II), we tested the binary response variable (deformed nymphs [DJ], live nymphs [LJ]) as a function of the predictors (i-iv). Third, using a Manova model (Type II), we tested the Differentially Expressed Genes (DEGs) of the aforementioned HSP genes (binary response variable [HSP70, HSP83]) as a function of the predictors (i-iii). The models were parsimonious as the non-significant predictors were removed. Each model was followed by a posthoc TUKEY test of pairwise comparisons; *emmeans* package [42].

### RNA extraction, quantification, and quality assessment

Total RNA was extracted using RNeasy Mini Kit (Qiagen©, UK) according to the manufacturer’s protocol including an on-column DNase digestion step. The DNase digestion was done using an RNase-free DNase set (Qiagen©, UK) according to the manufacturer’s protocol. Extracted RNA was quantified by Qubit® 3.0 Fluorometer using RNA Broad-Range (BR) Assay Kits (Invitrogen, Life Technologies©) according to the manufacturer’s instructions. RNA purity was checked using a Nanodrop spectrometer (Thermo Scientific©, UK) loading 1 μL eluted RNA sample and the 260/280 and 260/230 absorbance ratios were recorded. RNA integrity was assessed by running the samples in 1.5% agarose gel and high-quality DNA-free RNA samples were used for gene expression analysis using Quantitative reverse transcriptase PCR (qPCR).

### qPCR

RNA is reverse transcribed into complementary DNA (cDNA) using the QuantiTect Reverse Transcription kit (Qiagen©, UK) according to the manufacturer’s instructions. A geNorm analysis was conducted to determine suitable reference genes, SDHB and NADH, from a set of commonly used pea aphid reference genes, (S1 Table), that exhibit stability across experimental conditions (treatment group, genotype). The reaction volume in each well for the qPCR was 25 μL, comprising of 12.5 μL Quantifast SYBR green PCR master mix (Qiagen©, UK), 2.5 μL (10μM) forward primer, 2.5 μL (10μM) reverse primer, and 7.5 μL sample cDNA diluted 1:50 in nuclease-free water. After mixing the reaction mixtures by flicking, samples were picofuged and were run on an AriaMx qPCR machine (Agilent©, USA) on the following protocol: 1 cycle of activation of HotStar Taq Plus DNA polymerase at 95°C for 5min; 40 cycles of cDNA strand dissociation at 95°C for 10sec followed by primer annealing and extension at 60°C for 30sec. Additionally, a melt curve analysis was run for each plate with an initial melt step at 95°C for 1 min, dropping to 55°C for 30sec, followed by a 0.5°C interval/s incremental increase from 55°C to 95°C and the cycle threshold (Ct) value was determined. All samples were run in duplicates.

All unknown sample template values were interpolated from an eight-point 4-fold serial dilution standard curve, run in each plate, constructed from pooled cDNA. The selection of candidate genes (HSP83 and HSP70) was made based on their function in stress response, development, and embryogenesis. Primers for these genes were designed by PrimerBlast (NCBI), (S2 Table). The reaction mixtures for qPCR were prepared and run on an AriaMx qPCR machine following the same protocol, as mentioned above. All standards and samples were run in duplicates. After the data collection, mean Ct values of duplicates were used to estimate the values of samples from the standard curve. To generate values of relative gene expression, sample values were normalised to reference gene values, obtained from the same cDNA dilution. To confirm the amplification of a single qPCR product, a differential melt curve analysis was used.

### FTIR

To establish the differences in chemical compositions in response to stress, FTIR spectra were obtained and compared between samples of the aphid clones reared at 22°C or 27°C, at the end of the experiment. Aphid mothers were individually sampled in separate tubes and three replicates per condition were used for the FTIR measurement. The samples were analysed using a Vertex 70/70v FTIR spectrometer with Opus ver. 7.5 (Bruker, Germany) at the Department of Chemical Engineering, UoM. Each aphid was placed on the device crystal and squashed gently immediately after removing it from a dry ice box to avoid any degradation. Background and aphid sample spectra were recorded at 4 cm^−1^ resolution and 400 – 4000 cm^−1^ range from 32 scans. Averages of three replicates spectra across conditions and aphid clones were analysed and compared. The comparison was made for functional groups/wavelengths related to the metabolism of carbohydrates (1000 cm^−1^ – 1300 cm^−1^), proteins (1400 cm^−1^ – 1800 cm^−1^), and lipids (2700 cm^−1^ – 3000 cm^−1^) [24,43].

## Results

### Aphid survival and production of deformed nymphs

The numbers of the survived mothers that developed from early instars, during 14 days, differed significantly across aphid clones (F_(1,58)_ = 7.39, P = 0.009). Survival under the thermal optimum (control) was slightly higher than under thermal stress. More N116 mothers survived in both conditions when compared to N127 mothers, as mortality was higher under thermal stress in N127, (Fig 1).

**Fig 1.**
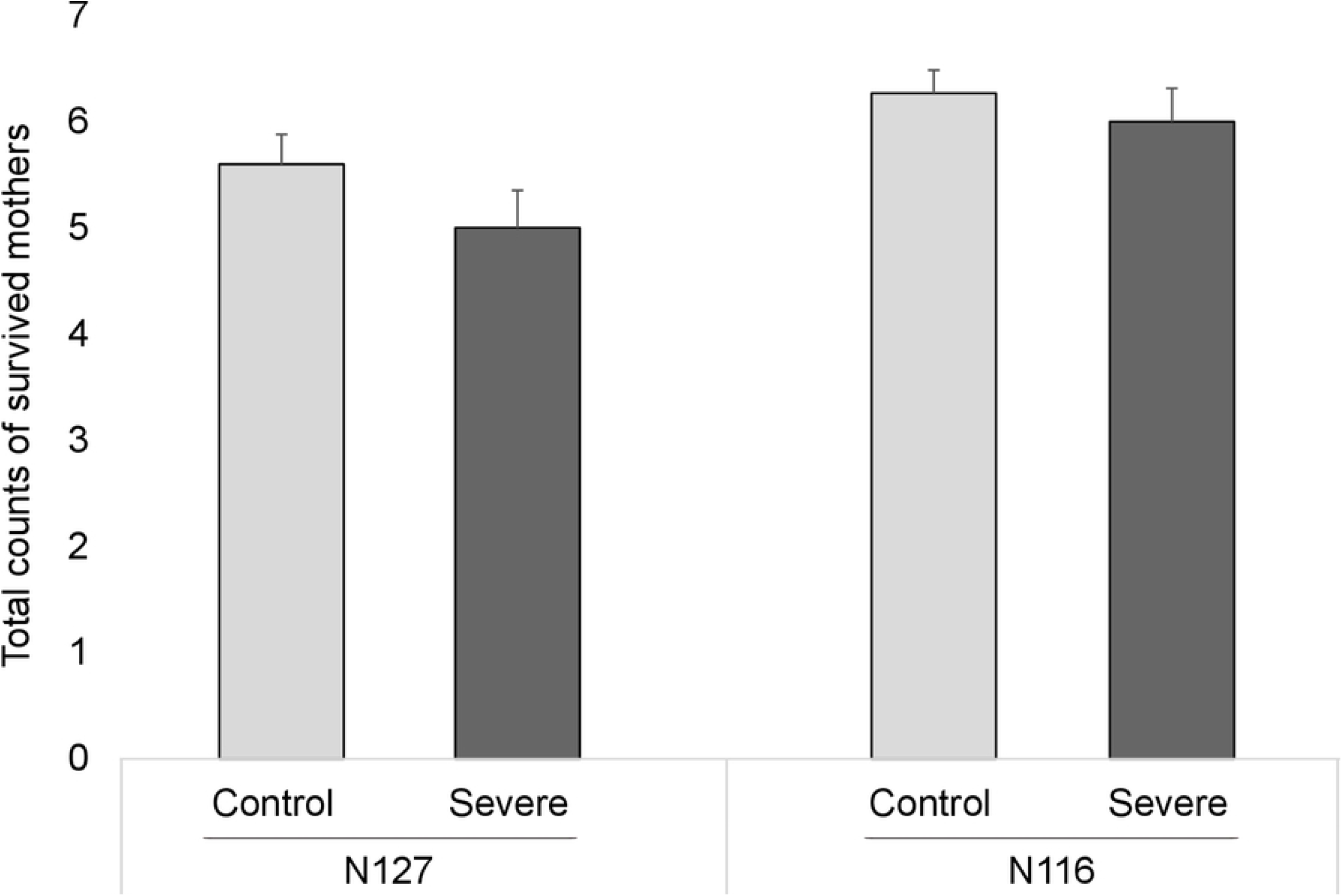
Aphid survival. The bars display the means (±SE) of the count of survived mothers at the end of the experiment on day 14, following their development from early instars. There were 60 enclosures in total including 2 aphid clones (N116 and N127) X 2 thermal conditions (control, 22°C or severe, 27°C) X 15 replicates.

Surviving mothers of the control produced live and healthy nymphs with functional appendages (Fig 2 [Panels F and G]). In contrast, subject to heatwave (thermal stress), surviving mothers produced striking scores of premature nymphs plus a few seemingly live but weak nymphs (Fig 2, [Panels A-E]). The majority of the nymphs produced by N127 were with malfunctioned appendages (late embryonic stage); appendage-less dead nymphs (early embryonic stage) were also detected (Fig 2). However, all the deformed progeny of N116 were appendage-less dead nymphs from an early embryonic stage.

**Fig 2.**
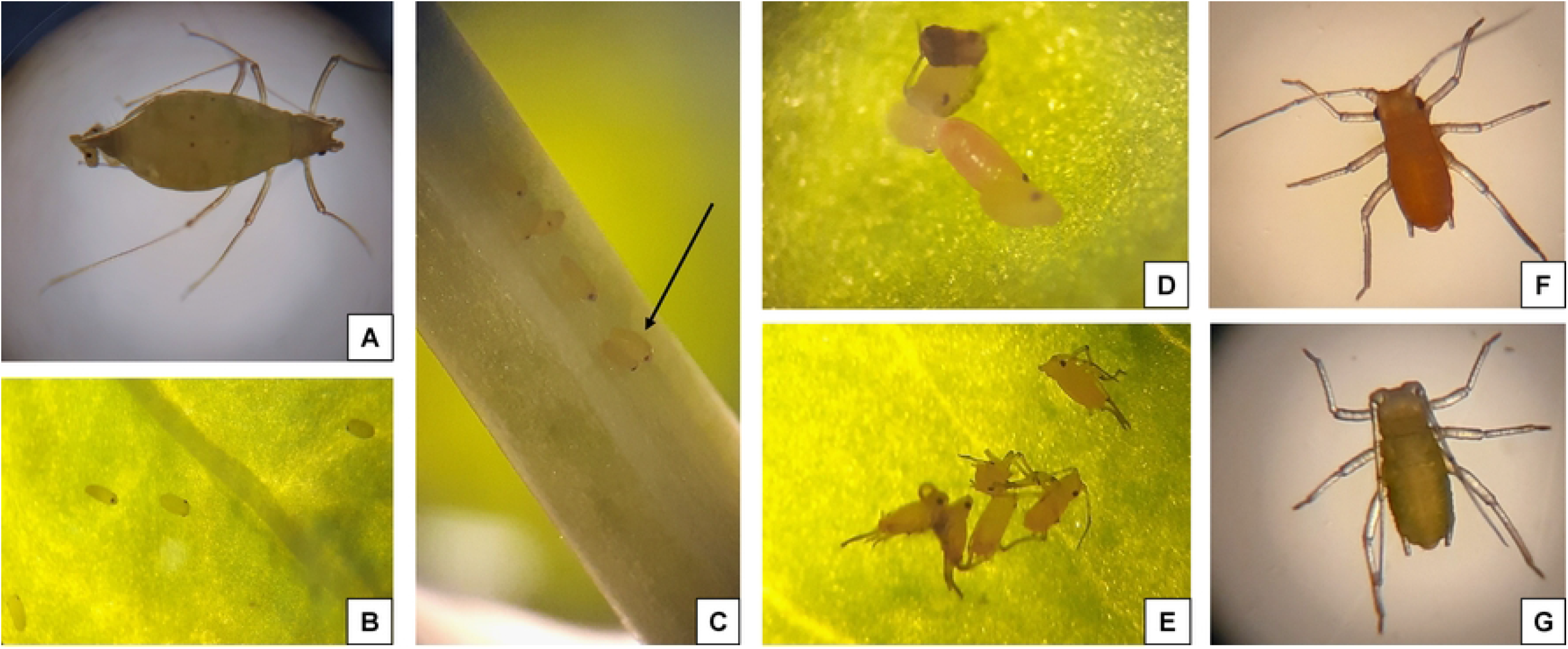
Premature/deformed nymph production in pea aphid clones under thermal stress. The aphids of both clones that developed under 27°C (heatwave) produced premature deformed nymphs and thrifty dull nymphs that generally appeared normal. Panel (A): an aphid mother (N116) producing a premature deformed nymph; Panel (B): some nymphs were born in early embryonic stages without any appendages (N116); Panel (C): An arrow pointing at a deformed nymph (N116) laid on the stem and lacking appendages; Panel (D): a deformed nymph (N127) with no appendages; Panel (E): some nymphs (N127) were born at their late embryonic stage but with folded (malfunctioning) appendages. In contrast, Panels (F) and (G) show close-ups of live healthy nymphs (N127 and N116, respectively) that were produced by aphids that developed under favourable temperate conditions (thermal optimum of 22°C).

Only healthy nymphs were produced by both aphid clones in the control. Conversely, development under thermal stress not only resulted in a sharp decline in aphid population size but also the deformities documented in (Fig 2). For clone N127, the average number of live healthy nymphs in the control was ~26 times that recorded in the stressful environment. A similar pattern, yet more pronounced, was observed for the N116 clone, as the average count of live healthy nymphs was ~34 times that of the control compared to thermal stress. Furthermore, the average number of healthy nymphs produced in the control by the N116 mothers was ~2 times what the N127 mothers produced, indicating that the N116 clone was more prolific than N127 independent of any treatment. However, under thermal stress, the numbers of live and deformed nymphs were slightly higher in N116 than in N127 (Fig 3). Again, interestingly, there were no deformed or premature nymphs under the favourable hospitable conditions of the control (Fig 3).

**Fig 3.**
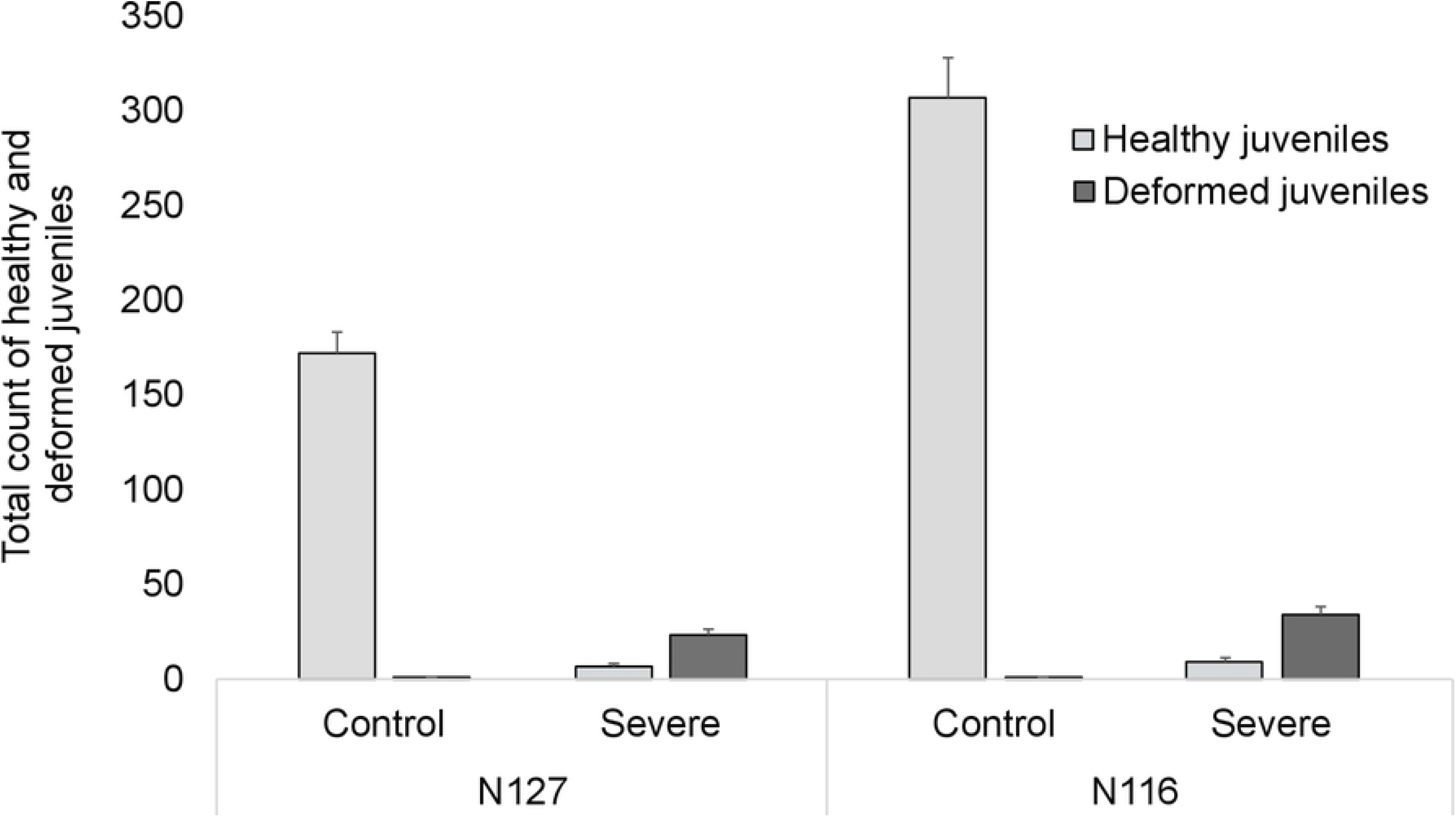
Total numbers of nymphs. The bars display the means (±SE) of the count of nymphs at the end of the experiment on day 14. Healthy (light grey) and deformed (dark grey) nymphs were produced by aphid mothers that developed and survived the heatwave. There were 60 enclosures in total including 2 aphid clones (N116 and N127) X 2 thermal conditions (control, 22°C or severe, 27°C) X 15 replicates.

The inferential stats revealed that thermal stress, aphid clone, and their interaction had highly significant effects on the numbers of both live and deformed nymphs (F_(1,51)_ = 234.76, P < 0.0001), (F_(1,51)_ = 19.04, P < 0.0001), and (F_(1,51)_ = 18.31, P < 0.0001), respectively. Furthermore, TDW and the interaction (TDW X Thermal Stress X Aphid clone) had significant effects (F_(1,51)_ = 4.02, P = 0.024) and (F_(1,52)_ = 3.71, P = 0.031), respectively. Under thermal stress, TDW decreased slightly for N127-infested plants, while it considerably increased for the N116-infested plants, with the margin of difference in TDW between the thermal conditions being higher in the case of N116; see (S3 Table) for full model details, and (S4 Table) for TDWs. The posthoc TUKEY test revealed only the following pairwise comparisons as significant (N127–control *vs*. N127–severe, P < 0.0001), (N116–control *vs*. N116–severe, P < 0.0001), and (N116–control *vs*. N127–control, P < 0.0001).

### HSP responses

HSP83 showed lower expression in the severe environment for both clones but the difference in comparison with the control was more apparent in N127, although the levels of expression were higher in N116 (Fig 4). Moreover, a higher HSP70 expression was detected in the severe environment and that was more prominent in N127 than in N116. The inferential stats revealed that thermal stress (F_(1,7)_ = 11.59, P = 0.006) and the interaction between aphid clone and thermal stress regime (F_(1,7)_ = 8.69, P = 0.013) had significant effects on the expression of both heat shock protein genes (HSP70 and HSP83). The effect of aphid clone was marginally significant (F_(1,7)_ = 3.76, P = 0.078). The posthoc TUKEY test only revealed the pairwise comparison (N127–control *vs*. N127–severe, P = 0.023) as significant; see (S5-S7 Tables) for an alternative analysis perspective. As such, qPCR data showed significant differential expression in HSP83 and HSP70 genes in N127 (but not in N116).

**Fig 4.**
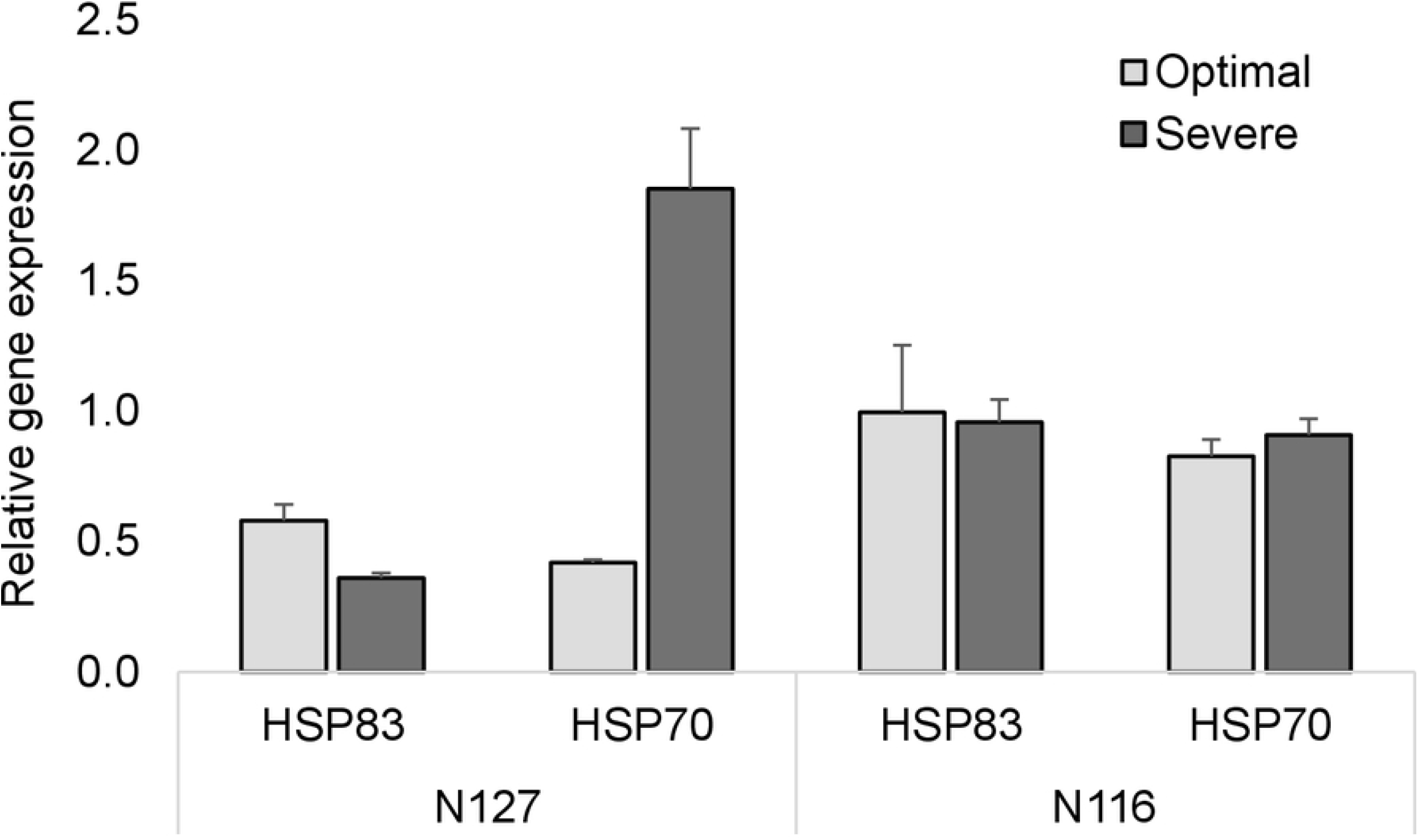
Relative gene expression (HSP83 and HSP70). The bars display the means (±SE) of relative gene expression of the aphid mothers in the control condition (light grey) and aphid mothers in the severe condition (dark grey) belonging to two aphid clones N116 and N127 at the end of the experiment. Gene expression was measured from three biological repeats consisting of pooled aphids from the respective genotypes across conditions, as specified in the Materials and methods.

### Metabolomic response

According to the FTIR spectra, lipids and fatty acids showed distinct higher intensities in the range (2849 – 2917 cm^−1^) under thermal stress. A less prominent difference between the thermal conditions was seen at the wavelength (2957 cm^−1^) (Figs 5 and 6). As for protein metabolism under thermal stress, distinct lower intensities within the range (1520 – 1627 cm^−1^), whilst a higher intensity at (1736 cm^−1^) were detected compared to the control. The patterns observed for the metabolism of lipids and proteins were generally identical for both aphid clones with slightly higher spikes for protein metabolism in N127 (Figs 5 and 6). Moreover, in the carbohydrate region, N116 showed a negligible difference between the two thermal conditions within the range (999 – 1154 cm^−1^), but N127 had a tangible difference with higher intensities under thermal stress. However, comparative to the control, both clones displayed lower intensities at (1240 cm^−1^), (Figs 5 and 6).

**Fig 5.**
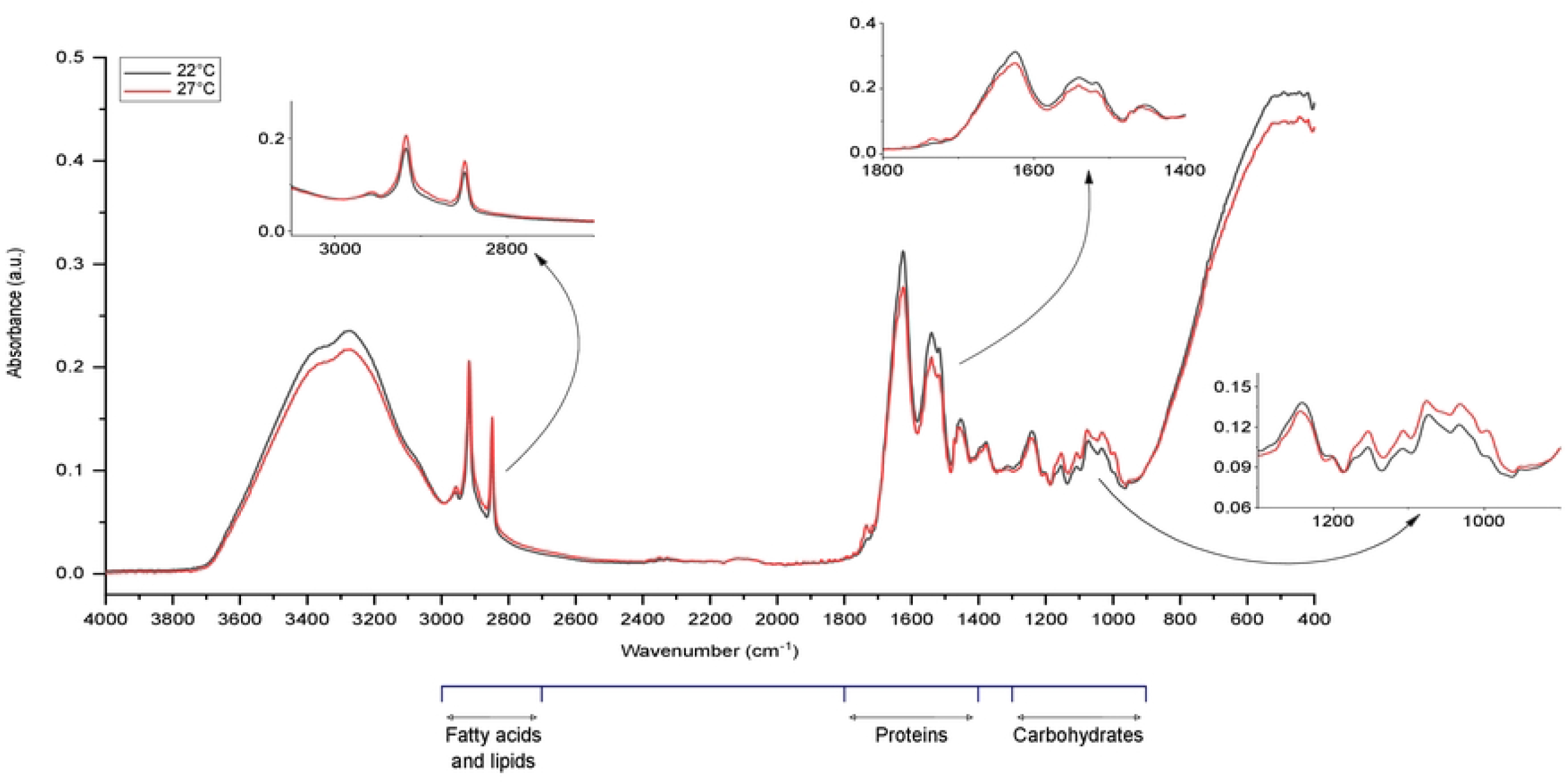
FTIR spectra of pink pea aphid (N127) in response to thermal stress. The red line represents the spectrum for the aphids of the N127 clone raised under heat stress (27°C), while the black line represents the spectrum for the aphids raised at a favourable temperature (thermal optimum of 22°C). Metabolic changes (carbohydrates, proteins, fatty acids and lipids) of the aphid body were compared. The main spectroscopic regions are in the wavelength range of 4000 – 400 cm^−1^.

**Fig 6.**
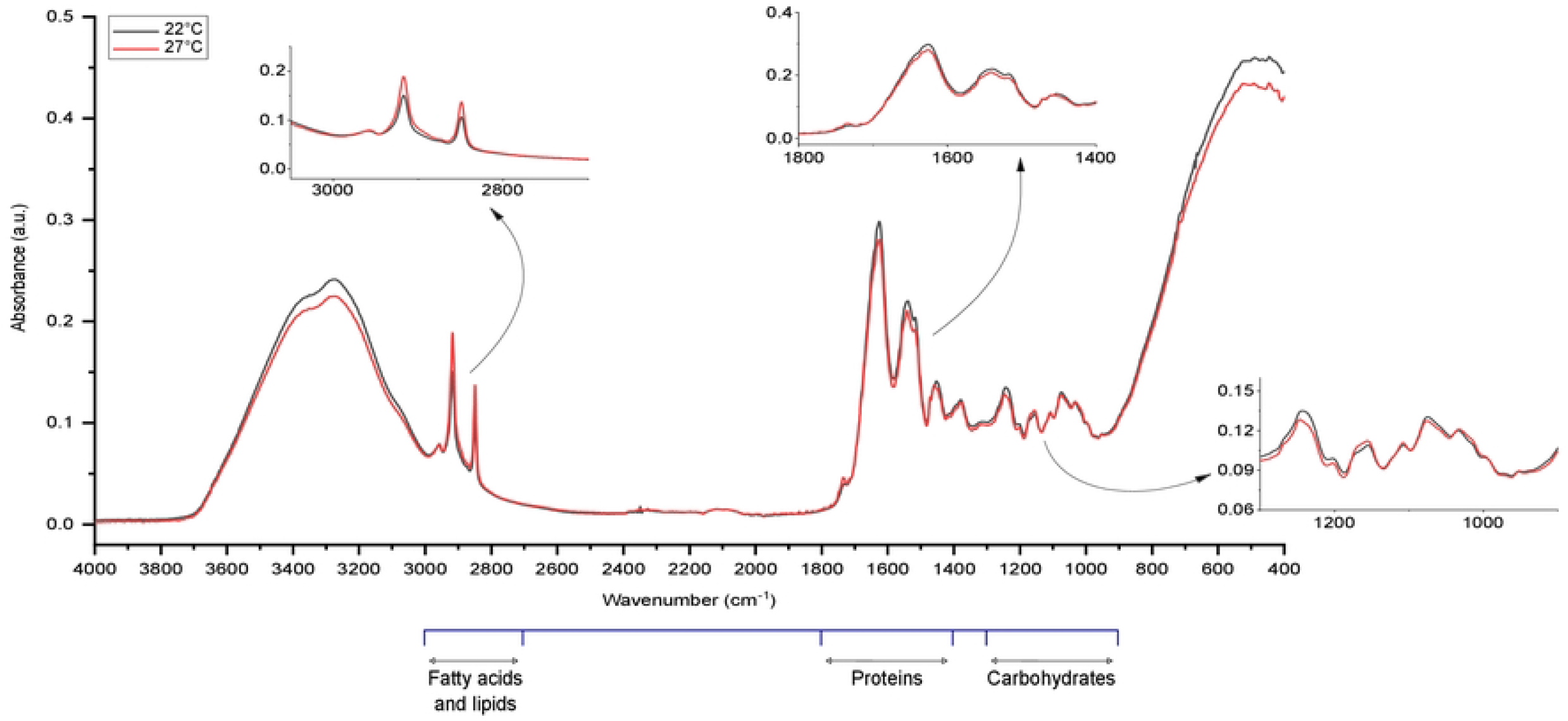
FTIR spectra of green pea aphid (N116) in response to thermal stress. The red line represents the spectrum for the aphids of the N127 clone raised under heat stress (27°C), while the black line represents the spectrum for the aphids raised at a favourable temperature (thermal optimum of 22°C). Metabolic changes (carbohydrates, proteins, fatty acids and lipids) of the aphid body were compared. The main spectroscopic regions are in the wavelength range of 4000 – 400 cm^−1^.

## Discussion

After 14-day exposure to thermal stress, the majority of aphid mothers of both clones survived but varyingly produced unexpected premature nymphs with deformities. These were either dead nymphs that lacked appendages or live deformed nymphs with folded malfunctioning appendages that died shortly. Whereas, the rest of the nymphs were weak and dull yet remained alive until the end of the experiment. We observed differential clone-specific expression of two HSP genes in the aphid mothers that developed, matured under stress, and survived compared to those reared in a thermal optimum. Additionally, certain metabolic changes were detected in response to thermal stress.

As for the survival of mothers, although the mortality rate in our study is less than that reported by Enders et al. (2015) [15], our findings highlight the detrimental impact of constant thermal stress on aphid survival in a context-dependent and clone-specific manner. The populations of the two clones dropped sharply whilst being exposed to the applied heatwave. This resonates with the findings of Chen et al. (2000) [44] on a reduced pea aphid fecundity at 25°C, and those of Dampc et al. (2021) [45] who found that exposure to 28°C affected the reproduction and longevity of rose aphid *Macrosiphum rosae* (Linnaeus) by altering plant-aphid interaction and their defence mechanism. Survival and nymph production in the severe condition were higher in N116 than in N127. That is attributable to a combination of factors including differences stemming from life-history traits [39], as well as stress responses on protein and metabolic levels as shown by our findings and discussed below. More importantly, the exposure of the mothers to a constant sub-lethal heatwave in the present work led to developmental anomalies in the offspring. This incomplete or unsuccessful embryogenesis is quite an unusual phenomenon, which to the best of our knowledge, has never been reported before ex-ovaries. It is worthy of note that to date, only artificial knockdown of the genes HSP83 [33] and Lysine Acetyltransferase p300/CBP (HAT) [34] in pea aphid resulted in the formation of premature nymphs with folded appendages.

We discuss our findings from different vantage points. Firstly, our results lend support to those of Shingleton et al. (2003) [46] who examined extreme developmental regulation and the occurrence of embryonic malformation during diapause. A higher rate of developmental abnormalities is detectable when the embryos develop at maintained perturbing high and/or fluctuating temperatures [46]. However, since diapause occurs at lower temperatures, it could be argued that aphids in our work experienced a state similar to quiescence that is usually associated with higher temperatures or abrupt and unexpected environmental change [47]. Given that the aphid clones here have been maintained in the lab for hundreds of generations where they never experienced heatwaves, the sudden change into unfavourable high sub-lethal temperature triggered a state similar to quiescence with a high risk of anomaly or physiological/developmental malfunction. The arrested/compromised development of some embryos [46], with a plausible state similar to quiescence, can be a compensatory mechanism that goes in line with interpreting fitness as a propensity [48] where only the fittest survive [48,49].

Secondly, aphids show rapid responses to harsh conditions through instant preconditioning of developing offspring, a phenomenon of extreme phenotypic plasticity (polyphenism) [19,28,50]. A trade-off usually occurs where aphid mothers strive to balance between producing more energetically costly *alatae* to escape adversity and the default norm of producing *apterae* that are more fecund yet vulnerable to environmental risk [28,51]. In analogy, developing and maturing in a thermally challenging environment, most of the viviparous aphid mothers in the current work survived, but they produced a mix of dead, malformed, and few semi-healthy offspring. The sub-lethal heatwave partly compromised aphid’s ability to cope with (or acclimate to) the constant stress exposure, challenging the balance between survival, development, and reproduction [52], but see [14]. A trade-off between survival and development might have taken place in the parthenogenetic all-female aphid populations, with arguably a plausible conflict between mothers and some of their developing embryos as well as between embryos [53,54] against nutrient deficiency resulting from critical changes in the density of their endosymbiont communities as we discuss below. It would be worth trying to entertain the prospect of sacrificing a portion of the developing daughters for the advantage of maintaining homeostasis in the bodies of their mothers. Aphids display exemplary cost-sensitive altruism against aphidophagous insects [55–58], but little is known on altruistic reproductive plasticity under severe physical conditions. If selective pressure favours altruism in harsh environmental conditions, as against natural enemies, future work is required to investigate a possibility of an altruism-orchestrating green-beard effect plausibly brought about by a nexus of genes associated with severe heat stress [54,59–61]. Even so, the question remains whether the observed phenomenon is an induced bet-hedging compensatory mechanism [62] under a thermally pressing ecological crunch or just a mere heat injury [63,64].

Thirdly, functional synthesis, mobilising, and conversion of metabolites are essential in maintaining energy homeostasis, embryogenesis, and stress resistance [65–67]. However, exposure to high temperatures may alter the functionality of these processes and result in deformities and compromised fitness as our findings suggest. Thermal stress depleted HSP83 expression in N127 but not in N116, which receives support from Will et al. (2017) [33] who found that reduced/depleted levels of HSP83 might lead to premature nymph development in aphid ovaries. HSP83 is a member of the HSP90 family of molecular chaperones that function as a determinant of fitness under non-optimal thermal conditions [14]. By contrast, HSP70 expression, which is usually upregulated under severe conditions [14], was higher under thermal stress in comparison with the control, with the highest level seen in N127. HSP70 is heavily involved in stress responses on many levels including synthesis, transcription, and metabolism [68,69]; elevated levels of HSP70, despite being essential to lessen stress burden, are energetically costly as it is ATP-dependent [68]. As such, the involvement of HSP genes in maintaining homeostasis, as well as anti-stress responses, may result in additional energy consumption to accommodate extra demands on resources, and that in turn may incur a fitness cost. There is evidence that an increased expression of HSP genes under thermal stress may negatively affect the natural development of embryos [46], leading to abnormal development, as shown in *D. melanogaster* [14]. Overall, the embryonic and larval defects examined in the current work could be the product of side effects of an increased upregulation of HSP70 for protection and maintenance as an anti-stress response, but our findings suggest that this may be clone-specific. The reported defects could also result from exhaustion or downregulation of HSP83 such that the expression of this HSP gene was insufficient to minimise embryonic anomalies and heat injuries in the exposed aphids.

Further, it is possible that the aphids in our study underwent a war of attrition between allocating resources for normal life functionality (including homeostasis) and reducing heat injuries [15,70] that otherwise can be disruptive or detrimental [7,71]. The constant heatwave could have also aggravated the thermal challenge not only by impacting population dynamics [52] but also possibly by altering osmoregulation and water content in the insect body [72–75], as well as energy reserves [1,76]. There was a distinct rise in metabolomic fingerprints of lipids and fatty acids following exposure to thermal stress, which was consistent across clones. Lower reproductive rates under thermal stress may protect the accumulation and utilisation of saturated fatty acids and that can aid in alleviating the energetically costly effects of stress [77]. This is well documented under lower temperatures, *e.g*., diapause [78], but consistent with the findings provided by Klepsatel et al. (2019) [79] on *D. melanogaster*, the described lipid metabolic response may be universal under low or high temperatures. Body fat is vital for functional metabolism and ovaries, and normal embryogenesis [64,78,80]. Thus, metabolic changes in body fat may reflect a possible adaptive stress response since insects must regulate the content and metabolism of their body fat [78,81,82] to offset stress-associated deficiencies [64,83,84]. All in all, mobilising and converting the storage of lipids and fatty acids can aid in counterbalancing the impacts of stress [85], corroborating our findings of active lipid metabolism in response to exposure to thermal disturbance. In a different light, conforming to the lipoid liberation theory [63,86], the constant exposure to non-lethal thermal stress in our study might have caused heat coagulation rather than protein denaturation that underlined embryonic deformities [63,86]. The coagulation might have compromised the essential roles of lipids in the haemolymph necessary for utilising energy, information signalling, and hence normal embryogenesis. Interestingly, thermal stress led to lower metabolic activity in the range (3000 cm^−1^ – 3600 cm^−1^), suggesting small changes in amides A and B regions (protein conformation) and membrane lipids [43].

Last but not least, aphid sensitivity to rising temperature and thermotolerance are associated with, or dependent on, their endosymbiont communities [12,44,87,88]. The density of the obligatory endosymbiont *Buchnera aphidicola* may vary according to aphid levels of maturity and exposure to thermal stress [89]. Insufficient availability of amino acids in the aphid diet due to lower densities of *B. aphidicola* in the thermally exposed hosts [12] could have also contributed to the production of deformed offspring in the current study. We note that N116 [58] and N127 harbour a range of facultative endosymbionts, with significantly lower titers in N127 (unpublished data). This difference in the titre may contribute to mitigating the negative effects of stressors on aphids [90] and also influence clone-specific gene expression patterns.

## Conclusions

The aphid clones in this study showed a continuum between ontogenetic plasticity, physiological change to resist stress, and failure to reproduce normally, although their populations did not go extinct. It was costly for the aphid mothers to survive in a constantly stressful thermal environment, as they incurred a fitness cost. Dead or premature and deformed offspring were produced due to compromised embryonic development. Aphid responses were shaped by the intertwined effects of endosymbiont density, life history, metabolic changes, differential expression of HSP genes, and a possible conflict between the survival of parthenogenetic mothers and the production of their offspring. This work casts light on a surprising response to atypical sub-lethal thermal stress and suggests a possibility for reproductive altruism, as it does for describing a new vulnerability of this important model organism and pest to heatwaves. Further research is necessary to examine whether this is unique to pea aphid or whether it may be more of a universal aphid anti-stress response.

## Acknowledgements

We are thankful to Ms. Gemma Chapman and Dr. Abdullatif Alfutimie of the Chemical Engineering Department (CE), UoM, for their support at the FTIR facility therein.

## Supporting information

**S1 Fig. Experimental design.** Seven first instars of each aphid clone (N127 [pink] or N116 [green]) were introduced to 2-week-old plants of one cultivar of fava bean grown and maintained at 22°C (thermal optimum) or 27°C (constant heatwave as thermal stress). Aphids were reared in the respective conditions for 14 days where they developed and reproduced. On day 15, aphid total numbers per enclosure (n=60) were counted and samples were taken for further analyses. Plants were washed at the end of the experiment and dried out in an oven at 55°C for two nights.

**S1 Table. List of reference genes tested to check expression stability.** The table shows five reference genes that were randomly selected from a list of ten common genes in pea aphid under different temperature conditions where the genes showed higher stability. See main text Methods for further details.

**S2 Table. Details of two candidate genes (HSP70 and HSP83) used in qPCR.** The table shows the name, function, primer sequence, and product size of the HSP genes that were selected from RNAseq results for validation. The differential expression levels of these genes were tested in two pea aphid clones (N127 and N116) under thermal stress (27°C) compared to thermal optimum (22°C). See main text Methods for further details.

**S3 Table. Analysis of the total number of nymphs (live and deformed).** The table shows the outcome of the Manova (Type II) model testing the total number of nymphs produced by the survived mothers. The predictors were (i) Thermal stress (two levels: 22°C [thermal optimum, control], 27°C [thermal stress]); (ii) Aphid clone (N116 [green], N127 [pink]), (iii) plant total dry weight (TDW) as a covariate, and (iv) the interactions of these predictors. The binary response variable was (deformed nymphs [DJ], and live nymphs [LJ]). See main text Methods for further details. Significant results are shown in bold. TS = thermal stress, Aphid C = Aphid clone.

**S4 Table. Total plant dry weight (TDW).** Means (± SE), standard deviation (SD), and relative standard deviation (RSD) of the total dry weight of the fava bean host plant. There were 60 enclosures in total including 2 aphid clones X 2 thermal conditions [control (favourable conditions at 22°C or stress at 27°C)] X 15 replicates. See main text Methods for further details.

**S5 Table. Analysis of HSP70 expression.** The table shows the outcome of the Anova (Type II) model testing changes in the expression of the HSP70 gene in two pea aphid clones (N127 and N116) under thermal stress (27°C) compared to thermal optimum (22°C). See main text Methods for further details. Significant results are shown in bold.

**S6 Table. Posthoc test of HSP70 expression.** The table shows the outcome of the posthoc TUKEY test following the Anova (Type II) model testing changes in the expression of the HSP70 gene in two pea aphid clones (N127 and N116) under thermal stress (27°C) compared to thermal optimum (22°C). See main text Methods for further details. Only significant results are shown.

**S7 Table. Analysis of HSP83 expression.** The table shows the outcome of the Anova (Type II) model testing changes in the expression of the HSP83 gene in two pea aphid clones (N127 and N116) under thermal stress (27°C) compared to thermal optimum (22°C). See main text Methods for further details. Significant results are shown in bold.

## Declarations

### Authorship contribution statement

Hawa Jahan (HJ): Conceptualisation, methodology, investigation, formal analysis, data curation and visualisation, resources, writing (original draft, review, and editing), funding acquisition, co-produced the manuscript with MSK.

Mouhammad Shadi Khudr (MSK): Conceptualisation, methodology, project administration, formal analysis and visualisation, writing (original draft, review, and editing), co-produced the manuscript with HJ.

Ali Arafeh (AA): Formal analysis, visualisation, writing (review).

Reinmar Hager (RH): Methodology, writing (review, and editing), resources, project administration, supervision.

### Data availability statement

The data underlying the results presented in the study are available from the [Figshare] repository via the following data link: [https://figshare.com/s/100808000e8d70fe036a].

## References

1. Ghaedi B, Andrew N. The physiological consequences of varied heat exposure events in adult *Myzus persicae*: a single prolonged exposure compared to repeated shorter exposures. PeerJ 4, p.e2290. 2016. doi: 10.7717/peerj.2290

2. Clarke B, Otto F, Stuart-Smith R, Harrington L. Extreme weather impacts of climate change: an attribution perspective. Environ. Res.: Climate. 2022; 1, 012001. doi: 10.1088/2752-5295/ac6e7d

3. Rouquette P. Climate change leading to earlier and earlier heatwaves, scientists say. France24. 2022. Available from: https://www.france24.com/en/europe/20220618-climate-change-leading-to-earlier-and-earlier-heatwaves

4. WHO. Climate change is increasing the risk of heatwaves: preparing for a warm and dry summer in the European Region. World Health Organization (WHO). 2022. Available from: https://www.who.int/europe/news/item/17-05-2022-climate-change-is-increasing-the-risk-of-heatwaves--preparing-for-a-warm-and-dry-summer-in-the-european-region

5. WMO. Temperature records tumble in early, intense heatwave. World Meteorological Organization (WMO). 2022. Available from: https://public.wmo.int/en/media/news/temperature-records-tumble-early-intense-heatwave

6. Ahn J, Cho J, Kim J, Seo B. Thermal Effects on the population parameters and growth of *Acyrthosiphon pisum* (Harris) (Hemiptera: Aphididae). Insects. 2020; 11, 481. doi: 10.3390/insects11080481

7. Sales K, Vasudeva R, Gage M. Fertility and mortality impacts of thermal stress from experimental heatwaves on different life stages and their recovery in a model insect. R. Soc. Open Sci. 2021; 8. doi: 10.1098/rsos.201717

8. Jørgensen K, Sørensen J, Bundgaard J. Heat tolerance and the effect of mild heat stress on reproductive characters in *Drosophila buzzatii* males. J. Therm. Biol. 2006; 31: 280–286. doi: 10.1016/j.jtherbio.2005.11.026

9. Zhang W, Rudolf V, Ma C. Stage-specific heat effects: timing and duration of heat waves alter demographic rates of a global insect pest. Oecologia. 2015; 179: 947–957. doi: 10.1007/s00442-015-3409-0

10. Chen H, et al. Effect of short-term high-temperature exposure on the life history parameters of *Ophraella communa*. Sci. Rep. 2018; 8. doi: 10.1038/s41598-018-32262-z

11. Stacey D, Fellowes M. Influence of temperature on pea aphid *Acyrthosiphon pisum* (Hemiptera: Aphididae) resistance to natural enemy attack. Bull. Entomol. Res. 2002; 92: 351–357. doi: 10.1079/ber2002173

12. Zhang B, Leonard S, Li Y, Moran N. Obligate bacterial endosymbionts limit thermal tolerance of insect host species. PNAS. 2019; 116: 24712–24718. doi: 10.1073/pnas.1915307116

13. Hullé M, Cœur d’Acier A, Bankhead-Dronnet S, Harrington R. Aphids in the face of global changes. C. R. Biol. 2010; 333: 497–503. doi: 10.1016/j.crvi.2010.03.005

14. Sørensen J, Kristensen T, Loeschcke V. The evolutionary and ecological role of heat shock proteins. Ecol. Lett. 2003; 6: 1025–1037. doi: 10.1046/j.1461-0248.2003.00528.x

15. Enders L, et al. Abiotic and biotic stressors causing equivalent mortality induce highly variable transcriptional responses in the soybean aphid. Genes Genomes Genet. 2015; 5: 261–270. doi: 10.1534/g3.114.015149

16. Van Emden H F, Harrington R. Aphids as Crop Pests. CAB International, Wallingford; 2017. doi: 10.1079/9781780647098.0000

17. Srinivasan D, Brisson J. Aphids: A model for polyphenism and epigenetics. Genet. Res. Int. 2012; 2012: 1–12. doi: 10.1155/2012/431531

18. West-Eberhard M. 2005. Developmental plasticity and the origin of species differences. PNAS 102, 6543–6549.; doi: 10.1073/pnas.0501844102

19. Whitman D, Ananthakrishnan T N. Phenotypic plasticity of insects: Mechanisms and consequences. Science Publishers, Enfield, NH; 2009.

20. Caillaud M, Losey J. Genetics of color polymorphism in the pea aphid, *Acyrthosiphon pisum*. J. Insect Sci. 2010; 10: 1–13. doi: 10.1673/031.010.9501

21. Castañeda L, Figueroa C, Bacigalupe L, Nespolo R. Effects of wing polyphenism, aphid genotype and host plant chemistry on energy metabolism of the grain aphid, *Sitobion avenae*. J. Insect Physiol. 2010; 56: 1920–1924. doi: 10.1016/j.jinsphys.2010.08.015

22. Hu L, Gui W, Chen B, Chen L. Transcriptome profiling of maternal stress-induced wing dimorphism in pea aphids. Ecol. Evol. 2019; 9: 11848–11862. doi: 10.1002/ece3.5692

23. Wang X, Chen Z, Feng Z, Zhu J, Zhang Y, Liu T. Starvation stress causes body color change and pigment degradation in *Acyrthosiphon pisum*. Front. Physiol. 2019; 10. doi: 10.3389/fphys.2019.00197

24. Durak R, Depciuch J, Kapusta I, Kisała K, Durak T. Changes in chemical composition and accumulation of cryoprotectants as the adaptation of anholocyclic aphid *Cinara tujafilina* to overwintering. Int. J. Mol. Sci. 2021 Jan 6;22(2):511. doi: 10.3390/ijms22020511

25. Tougeron K, van Baaren J, Nordin D, Dumonceaux T, Wist T. Body-color plasticity of the English grain aphid in response to light in both laboratory and field conditions. Evol. Ecol. 2021; 35: 147–162. doi: 10.1007/s10682-020-10088-4

26. Jeffs C, Leather S. Effects of extreme, fluctuating temperature events on life history traits of the grain aphid, *Sitobion avenae*. Entomol. Exp. Appl. 2014; 150: 240–249. doi: 10.1111/eea.12160

27. Peng X. et al. Effects of variable maternal temperature on offspring development and reproduction of *Rhopalosiphum padi*, a serious global pest of wheat. Ecol. Entomol. 2019; 45: 269–277. doi: 10.1111/een.12796

28. Stadler B. Adaptive allocation of resources and life-history trade-offs in aphids relative to plant quality. Oecologia. 1995; 102: 246–254. Available from: http://www.jstor.org/stable/4220954

29. Laughton A, Fan M, Gerardo N. The Combined effects of bacterial symbionts and aging on life history traits in the pea aphid, *Acyrthosiphon pisum*. Appl. Environ. Microbiol. 2014; 80: 470–477. doi: 10.1128/aem.02657-13

30. Karunanithi S, Brown I. Heat shock response and homeostatic plasticity. Front. Cell. Neurosci. 2015; 9. doi: 10.3389/fncel.2015.00068

31. Goodman R H, Smolik S. CBP/p300 in cell growth, transformation, and development. Genes Dev. 2000; 14: 1553–1577. doi: 10.1101/gad.14.13.1553

32. Roberts S P, Feder M E. Natural hyperthermia and expression of the heat shock protein Hsp70 affect developmental abnormalities in *Drosophila melanogaster*. Oecologia. 1999; 121: 323–329. Available from: http://www.jstor.org/stable/4222473

33. Will T, Schmidtberg H, Skaljac M, Vilcinskas A. Heat shock protein 83 plays pleiotropic roles in embryogenesis, longevity, and fecundity of the pea aphid *Acyrthosiphon pisum*. Dev. Genes Evol. 2017; 227: 1–9. doi: 10.1007/s00427-016-0564-1

34. Kirfel P, Vilcinskas A, Skaljac M. Lysine acetyltransferase p300/cbp plays an important role in reproduction, embryogenesis and longevity of the pea aphid *Acyrthosiphon pisum*. Insects. 2020; 11: 265. doi: 10.3390/insects11050265

35. Allwood J, et al. Plant metabolomics, its potential for systems biology research. Meth. Enzymol. 2011; 500: 299–336. doi: 10.1016/B978-0-12-385118-5.00016-5

36. Lankadurai B, Nagato E and Simpson M. Environmental metabolomics: an emerging approach to study organism responses to environmental stressors. Environ. Rev. 2013; 21: 180–205. doi: 10.1139/er-2013-0011

37. Machovič V, et al. Analysis of European honeybee (*Apis mellifera*) wings using ATR-FTIR, Raman spectroscopy: A pilot study. Sci. Agric. Bohem. 2017; 48: 22–29. doi: 10.1515/sab-2017-0004

38. Ammar E, Alessandro R, Hall D. Ultrastructural and chemical studies on waxy secretions and wax-producing structures on the integument of the woolly oak aphid *Stegophylla brevirostris* Quednau (Hemiptera: Aphididae). J. Microsc. Ultrastruct. 2013; 1: p.43. doi: 10.1016/j.jmau.2013.05.001

39. Kanvil S, Powell G, Turnbull C. Pea aphid biotype performance on diverse Medicago host genotypes indicates highly specific virulence and resistance functions. Bull. Entomol. Res. 2014; 104: 689–701. doi: 10.1017/s0007485314000443

40. R Core Team 2021. R: A language and environment for statistical computing. R Foundation for Statistical Computing, Vienna, Austria. https://www.R-project.org/

41. Fox J, Weisberg S. An R Companion to Applied Regression, Third edition. Sage, Thousand Oaks CA; 2019. Available from: https://socialsciences.mcmaster.ca/jfox/Books/Companion/

42. Lenth R V. emmeans: Estimated marginal means, aka LeastSquares Means. R package version 1.1.2018. Available from: https://CRAN.R-project.org/package=emmeans

43. Correia M, et al. FTIR Spectroscopy - a potential tool to identify metabolic changes in dementia patients. Journal of Alzheimers & Neurodegenerative Diseases. 2016; 2: 007. doi: 10.24966/AND-9608/100007

44. Chen D, Montllor C, Purcell A. Fitness effects of two facultative endosymbiotic bacteria on the pea aphid, *Acyrthosiphon pisum*, and the blue alfalfa aphid, A. kondoi. Entomol. Exp. Appl. 2000; 95: 315–323. doi: 10.1046/j.1570-7458.2000.00670.x

45. Dampc J, Mołoń M, Durak T, Durak, R. Changes in aphid—plant interactions under increased temperature. Biology. 2021; 10: 480. doi: 10.3390/biology10060480

46. Shingleton A W, Sisk G C, Stern D L. Diapause in the pea aphid (*Acyrthosiphon pisum*) is a slowing but not a cessation of development. BMC Dev. Biol. 2003; 3: 7. doi: 10.1186/1471-213x-3-7

47. Durak R, Dampc J, Dampc J, Bartoszewski S, Michalik A. Uninterrupted development of two aphid species belonging to Cinara genus during winter diapause. Insects. 2020; 11: 150. doi: 10.3390/insects11030150

48. Mills S K, Beatty J H. The propensity interpretation of fitness. Philos. Sci. 1979; 46: 263–286. Available from: http://www.jstor.org/stable/187048

49. Sober E. Conceptual issues in evolutionary biology. 3rd edition. MIT Press, UK; 2006.

50. Dixon A. Aphid ecology. Chapman and Hall, London, UK; 1998.

51. Zhang Y, Wu K, Wyckhuys K, Heimpel G. Trade-offs between flight and fecundity in the soybean aphid (Hemiptera: Aphididae). J. Econ. Entomol. 2009; 102: 133–138. doi: 10.1603/029.102.0119

52. Zhao F, Zhang W, Hoffmann A, Ma C. Night warming on hot days produces novel impacts on development, survival and reproduction in a small arthropod. J. Anim. Ecol. 2013; 83: 769–778. doi: 10.1111/1365-2656.12196

53. Haig D. Gestational drive and the green-bearded placenta. PNAS. 1996; 93: 6547–6551. doi: 10.1073/pnas.93.13.6547

54. Grafen A. Green beard as death warrant. Nature. 1998; 394: 521–522. doi: 10.1038/28948

55. McAllister M, Roitberg B. Adaptive suicidal behaviour in pea aphids. Nature, 1987; 328: 797–799. doi: 10.1038/328797b0

56. Khudr M S, Oldekop J, Shuker D, Preziosi R. Parasitoid wasps influence where aphids die via an interspecific indirect genetic effect. Biol. Lett. 2013; 9: 20121151. doi: 10.1098/rsbl.2012.1151

57. Khudr M S, Fliegner L, Buzhdygan O, Wurst S. Super-predation and intraguild interactions in a multi-predator-one-prey system alter the abundance and behaviour of green peach aphid (Hemiptera: Aphididae). Can. Entomol. 2020; 152: 200–223. doi: 10.4039/tce.2020.7

58. Purkiss S A, Khudr M S, Aguinaga O E, Hager R. Symbiont-conferred immunity interacts with effects of parasitoid genotype and intraguild predation to affect aphid immunity in a clone-specific fashion. BMC Ecol. Evol. 2022; 22: 33. doi: 10.1186/s12862-022-01991-1

59. Hamilton W. The genetical evolution of social behaviour. I. J. Theor. Biol. 1964; 7: 1–16. doi: 10.1016/0022-5193(64)90038-4

60. Dawkins R. The Selfish Gene. Oxford Univ. Press, New York; 1976.

61. Trubenová B, Hager R. Green beards in the light of indirect genetic effects. Ecol. Evol. 2019; 9: 9597–9608. doi: 10.1002/ece3.5484

62. Joschinski J, Bonte D. Transgenerational plasticity and bet-hedging: a framework for reaction norm evolution. Front. Ecol. Evol. 2020; 8. doi: 10.3389/fevo.2020.517183

63. Munson S. Some effects of storage at different temperatures on the lipids of the American roach and on the resistance of this insect to heat and to D D T. Ph.D. Thesis. University of Maryland. 1952. Available from: https://drum.lib.umd.edu/bitstream/handle/1903/17831/DP70500.pdf?sequence=1

64. Downer R G H, Matthews J R. Patterns of lipid distribution and utilisation in insects. Am. Zool. 1976; 16: 733–745. doi: 10.1093/icb/16.4.733

65. Fraga A et al. Glycogen and glucose metabolism are essential for early embryonic development of the red flour beetle *Tribolium castaneum*. PLoS ONE. 2013; 8: e65125. doi: 10.1371/journal.pone.0065125

66. Li Y, et al. Evaluation of the expression and function of the tre2-like and tre2 genes in ecdysis of *Harmonia axyridis*. Front. Physiol. 2019; 10. doi: 10.3389/fphys.2019.01371

67. Bretscher H, O’Connor M. The role of muscle in insect energy homeostasis. Front. Physiol. 2020; 11. doi: 10.3389/fphys.2020.580687

68. Clare D K, Saibil H R. ATP-driven molecular chaperone machines. Biopolymers. 2013; 99: 846–859. doi: 10.1002/bip.22361

69. Cai Z, Chen J, Cheng J, Lin T. Overexpression of three heat shock proteins protects *Monochamus alternatus* (Coleoptera: Cerambycidae) from thermal stress. J. Insect Sci. 2017; 17. doi: 10.1093/jisesa/iex082

70. Jurivich D, Zhou X. Stress: physiological. In: Birren J E, editor. Encyclopedia of Gerontology (2^nd^ edition). Elsevier; 2007. pp. 559–565. doi: 10.1016/b0-12-370870-2/00179-7

71. Perez R, Aron S. Adaptations to thermal stress in social insects: recent advances and future directions. Biol. Rev. 2020; 95: 1535–1553. doi: 10.1111/brv.12628

72. Karley A J, Ashford D A, Minto L M, Pritchard J, Douglas A E. The significance of gut sucrase activity for osmoregulation in the pea aphid, *Acyrthosiphon pisum*. J. Insect Physiol. 2005; 51: 1313–1319. doi: 10.1016/j.jinsphys.2005.08.001

73. Beyenbach K W. The plasticity of extracellular fluid homeostasis in insects. J. Exp. Biol. 2016; 219: 2596–607. doi: 10.1242/jeb.129650

74. Sadras V O, et al. Aphid resistance: An overlooked ecological dimension of nonstructural carbohydrates in cereals. Front. Plant Sci. 2020; 11: 937.; doi: 10.3389/fpls.2020.00937

75. Liu J, Wang C, Desneux N, Lu Y. 2021. Impact of temperature on survival rate, fecundity, and feeding behavior of two aphids, *Aphis gossypii* and *Acyrthosiphon gossypii*, when reared on cotton. Insects 12, 565.; doi: 10.3390/insects12060565

76. Chown S, Sørensen J, Terblanche J. Water loss in insects: An environmental change perspective. J. Insect Physiol. 2011; 57: 1070–1084. doi: 10.1016/j.jinsphys.2011.05.004

77. Hubhachen Z, Madden R, Dillwith J. Influence of rearing temperature on triacylglycerol storage in the pea aphid, *Acyrthosiphon pisum. Arch*. Insect Biochem. Physiol. 2018; 99: p.e21495. doi: 10.1002/arch.21495

78. Arrese E, Soulages J. Insect fat body: energy, metabolism, and regulation. Annu. Rev. Entomol. 2010; 55: 207–225. doi: 10.1146/annurev-ento-112408-085356

79. Klepsatel P, Wildridge D, Gáliková M. Temperature induces changes in *Drosophila* energy stores. Sci. Rep. 2019; 9: 5239. doi: 10.1038/s41598-019-41754-5

80. Ziegler R, Vanantwerpen R. Lipid uptake by insect oocytes. Insect Biochem. Mol. Biol. 2006; 36: 264–272. doi: 10.1016/j.ibmb.2006.01.014

81. Keeley L. Physiology and biochemistry of the fat body. In: Kerkut G A, Gilbert, L I, editors. Integument, Respiration and Circulation. Pergamon; 1985. pp.211–248. doi: 10.1016/b978-0-08-030804-3.50012-1

82. Toprak U, Hegedus D, Doğan C, Güney G. A journey into the world of insect lipid metabolism. Arch. Insect Biochem. Physiol. 2020: 104:e21682. doi: 10.1002/arch.21682

83. Balogh G, et al. Key role of lipids in heat stress management. FEBS Lett. 2013; 587: 1970–1980. doi: 10.1016/j.febslet.2013.05.016

84. Jarc E, Petan T. Lipid droplets and the management of cellular stress. Yale J. Biol. Med. 2019; 92: 435–452. Available from: https://www.ncbi.nlm.nih.gov/pmc/articles/PMC6747940/pdf/yjbm_92_3_435.pdf

85. Arrese E L, Canavoso L E, Jouni Z E, Pennington J E, Tsuchida K, Well MA. Lipid storage and mobilization in insects: current status and future directions. Insect Biochem. Mol. Biol. 2001; 31: 7–17. doi: 10.1016/s0965-1748(00)00102-8

86. Heber U, Santarius K. Cell death by cold and heat, and resistance to extreme temperatures. Mechanisms of hardening and dehardening. In: Precht H, Christophersen J, Hensel H, Larcher, editors. Temperature and Life. Springer-Verlag, Berlin/Heidelberg; 1973. pp. 232–263.

87. Dunbar H, Wilson A, Ferguson N, Moran N. Aphid thermal tolerance is governed by a point mutation in bacterial symbionts. PLoS Biol. 2007; 5, e96. doi: 10.1371/journal.pbio.0050096

88. Corbin C, Heyworth E, Ferrari J, Hurst G. Heritable symbionts in a world of varying temperature. Heredity. 2016; 118: 10–20. doi: 10.1038/hdy.2016.71

89. Patton M, Hansen A, Casteel C. The green peach aphid, *Myzus persicae*, transcriptome in response to a circulative, nonpropagative polerovirus, Potato leafroll virus. BioRxiv [Preprint]. 2021 bioRxiv *2021.04.15.440077*. 2021. doi: 10.1101/2021.04.15.440077

90. Heyworth E, Smee M, Ferrari J. Aphid facultative symbionts aid recovery of their obligate symbiont and their host after heat stress. Front. Ecol. Evol. 2020; 8. doi: 10.3389/fevo.2020.00056

